# Regulatory divergence in wound-responsive gene expression in domesticated and wild tomato

**DOI:** 10.1101/123992

**Authors:** Ming-Jung Liu, Koichi Sugimoto, Sahra Uygun, Nicholas Panchy, Michael S. Campbell, Mark Yandell, Gregg A. Howe, Shin-Han Shiu

## Abstract

**Background:** The evolution of *cis-* and *trans-*regulatory components of transcription is central to how stress response and tolerance differ across species. However, it remains largely unknown how divergence in TF binding specificity and *cis*-regulatory sites contribute to the divergence of stress-responsive gene expression between wild and domesticated species.

**Results:** Using tomato as model, we analyzed the transcriptional profile of wound-responsive genes in wild *Solanum pennellii* and domesticated *S. lycopersicum*. We found that extensive expression divergence of wound-responsive genes is associated with speciation. To assess the degree of trans-regulatory divergence between these two species, 342 and 267 putative *cis-*regulatory elements (pCREs) in *S. lycopersicum* and *S. pennellii*, respectively, were identified that were predictive of wound-induced gene expression. We found that 35-66% of pCREs were conserved across species, suggesting that the remaining proportion (34-65%) of pCREs are species specific. This finding indicates a substantially higher degree of trans-regulatory divergence between these two plant species, which diverged ∼3-7 million years ago, compared to that observed in mouse and human, which diverged ∼100 million years ago. In addition, differences in pCRE sites were significantly associated with differences in wound-responsive gene expression between wild and domesticated tomato orthologs, suggesting the presence of substantial *cis*-regulatory divergence.

**Conclusions:** Our study provides new insights into the mechanistic basis of how the transcriptional response to wounding is regulated and, importantly, the contribution of *cis-* and *trans-*regulatory components to variation in wound-responsive gene expression during species domestication.

## BACKGROUND

Natural or artificial selection on diverse phenotypes leads to adaptation and domestication [1, 2]. Studies of the regulatory mechanisms underlying phenotypic diversity suggest that the variation in gene expression at the transcriptional level is one of the major contributing factors [3, 4]. To determine when, where and at what level genes are transcribed, the transcription regulatory program has two major components. The first is *trans-acting* factors including DNA-binding transcription factors (TFs) [5, 6]. The second component is the *cis-*regulatory sites [5-7], which are typically ∼6-15 bp in length and located in close proximity to their target genes. A TF generally recognizes multiple, slightly different *cis*-regulatory sites (collectively referred to as a *cis*-regulatory element (CRE) representing the binding specificity of TFs) [7]. Thus, variation in gene expression may result from the differences in the binding specificity of TFs and the CREs to which TFs bind. Differences in the specific *cis*-regulatory sites may also contribute to expression divergence between orthologs. In cross-species studies, CREs have been shown to evolve much slower than *cis*-regulatory sites that have undergone extensive divergence [3, 4, 8, 9]. For example, CREs amongst the orthologous TFs from mouse and human are highly conserved [10, 11]. Similarly, by identifying sequence motifs (resembling CREs) from mouse and human DNase I footprints, 95% of the mouse motifs were similar to those from human [10]. Because CREs are distinct TF binding motifs, these findings of CRE conservation indicate a high degree of conservation in *trans* regulatory mechanisms. In contrast, only ∼20% of mouse DNase I footprints were co-localized with human footprints [10], suggesting that the *cis*-regulatory sites have diverged extensively across species. But it remains unclear to what extent the *trans*-acting factors and *cis*-regulatory sequences have diverged in plants.

The divergent phenotypes between domesticated and wild plant species are the result of the domestication process in response to human selection [2, 12-16]. Comparisons of transcriptome profiles between domesticated and wild maize, carrot, cotton and tomato species have revealed that the extensive changes of gene expression are associated with phenotypic differences between closely related wild-domesticated species pairs [17-20]. In addition, studies have shown that the divergence of *cis*-regulatory sites affects the transcript level of key regulators involved in tomato leaf shape and fruit size and shape, rice slenderness, barley spike density and maize flowering time [2, 20, 21]. Thus, variation in *cis*-regulatory sites can contribute to domestication traits. In addition to domestication traits impacting development and physiology, domesticated and wild species tend to have significant differences in stress tolerance traits. This may be caused by artificial selection for desirable traits, which creates bottlenecks due to the selection of a small subset of individuals in the population and leaves behind genes relevant to biotic/abiotic tolerance compared to their wild species [13, 22, 23]. As a result, the wild species preserve much of the genetic variation underlying stress tolerance mechanisms and are frequently used to introgress stress tolerance traits into their domesticated counterparts [16, 18, 24, 25].

To better understand the roles of **cis*-* and *trans*-regulatory divergence between species in stress tolerance traits, the domesticated and wild tomato species are good models because of significant differences in stress tolerance between these species [16, 18, 24]. Recent transcriptomic studies in the domesticated tomato, *S. lycopersicum*, and the wild species, *S. pennellii*, have revealed a number of differential expressed genes involved in stress response [18, 24]. These studies have identified transposable elements (TEs) as a source of variation in *cis*-regulatory sites and control of stress-induced gene expression [24, 26, 27]. Among the diverse signals triggering the response against biotic/abiotic stress, response to wounding in tomato is among the most well characterized because of their relevance to herbivory [28, 29]. Studies in the domesticated tomato species *S. lycopersicum*, for example, have revealed that wounding induces changes in gene expression that ultimately trigger a defense-signaling network for resistance to insect herbivory [30-32]. In addition, several studies have demonstrated that stress-responsive TFs such as bHLHs and bZIPs and their corresponding *cis*-regulatory site are involved in transcriptional regulation of wound-responsive gene expression in *S. lycopersicum* [33, 34]. Nonetheless, the identities of most CREs and their corresponding regulatory sites underlying stress tolerance regulation and divergence in tomato or most other plant species have not been comprehensively examined. It also remains unclear how wound-induced patterns of gene expression differ between domesticated and wild tomato species such as *S. pennellii*, and how divergence in TFs (which can be inferred based on CREs) and *cis*-regulatory sites contribute to wound response divergence between these species.

In this study we investigated the divergence in wound-responsive gene expression and its corresponding regulatory mechanisms between the wild species *S. pennellii* and the domesticated *S. lycopersicum*. Specifically, we asked: (1) to what extent the wound-responsive gene expression has diverged between *S. lycopersicum* and *S. pennellii*, (2) what the CREs are that regulate differentially expressed genes between wound-treated and control samples from each species and between species, (3) to what degree are CREs relevant to wound-responsive gene expression conserved across species, and (4) to what extent are differences in wound-induced transcriptional responses in these two tomato species attributed to divergence in *cis*-regulatory sites.

## RESULTS and DISCUSSION

### Temporal and spatial expression profiles of wound-responsive genes in two Solanum species

To globally examine how the effects of wounding on gene expression differ in *S. pennellii* and *S. lycopersicum*, leaves were wounded mechanically to trigger the response in damaged (local) and undamaged (systemic) tissues and at different time points after wounding. The local and systemic leaves at 0.5- and 2-hr time points post-treatment were collected for transcript profiling. For comparison, control leaf tissue was collected from unwounded plants. A gene is defined as wound-responsive if it is either significantly up or down-regulated (adjusted p<0.05, |log_2_(FC)|>2, FC: Fold Change) in a wounded sample compared to the unwounded control. To increase the stringency of our analysis, we choose a FC threshold of four fold instead of the conventional two fold to emphasize robust changes in gene expression. In both species, ∼1,000 genes were significantly up-regulated by wounding (wound-induced) in local leaves during both early (0.5-hr) and late (2-hr) time points (**Fig. 1A**). Interestingly, the pattern is very different for down-regulated genes whose expression is repressed by wounding. At the 0.5 hr time point of the local response, only 71 *S. lycopersicum* genes were down-regulated compared to 602 in *S. pennellii* (**Fig. 1A**). Similarly, there were still fewer down-regulated genes in *S. lycopersicum* at the 2-hr time point of the local response (211; left panel, **Fig. 1A**) compared to *S. pennellii* (625; right panel, **Fig. 1A**). Thus for the local response, the general pattern of wound-induced gene expression was similar between the two species, whereas wounding repressed gene expression to a greater extent in the wild species relative to domesticated tomato.

Far fewer genes were differentially expressed during the systemic wound response in both species, with *S. pennellii* having more systematically responsive genes than the cultivated species (249 in *S. lycopersicum* and 593 in *S. pennellii*) (**Fig. 1A**). Approximately 83% and 75% of these systemic wound-responsive genes in *S. lycopersicum* and *S. pennellii*, respectively, were a subset of the local wound-responsive genes. Consistent with previous microarray studies [32], our results based on transcriptome sequencing indicate that there are similar wound responses between the local and systemic leaf (**Fig. 1A**). Taken together, these findings show that in response to wounding, both species have extensive changes in gene expression programs, but the extent of gene expression repression is more prominent in *S. pennellii*.

**Figure 1.**
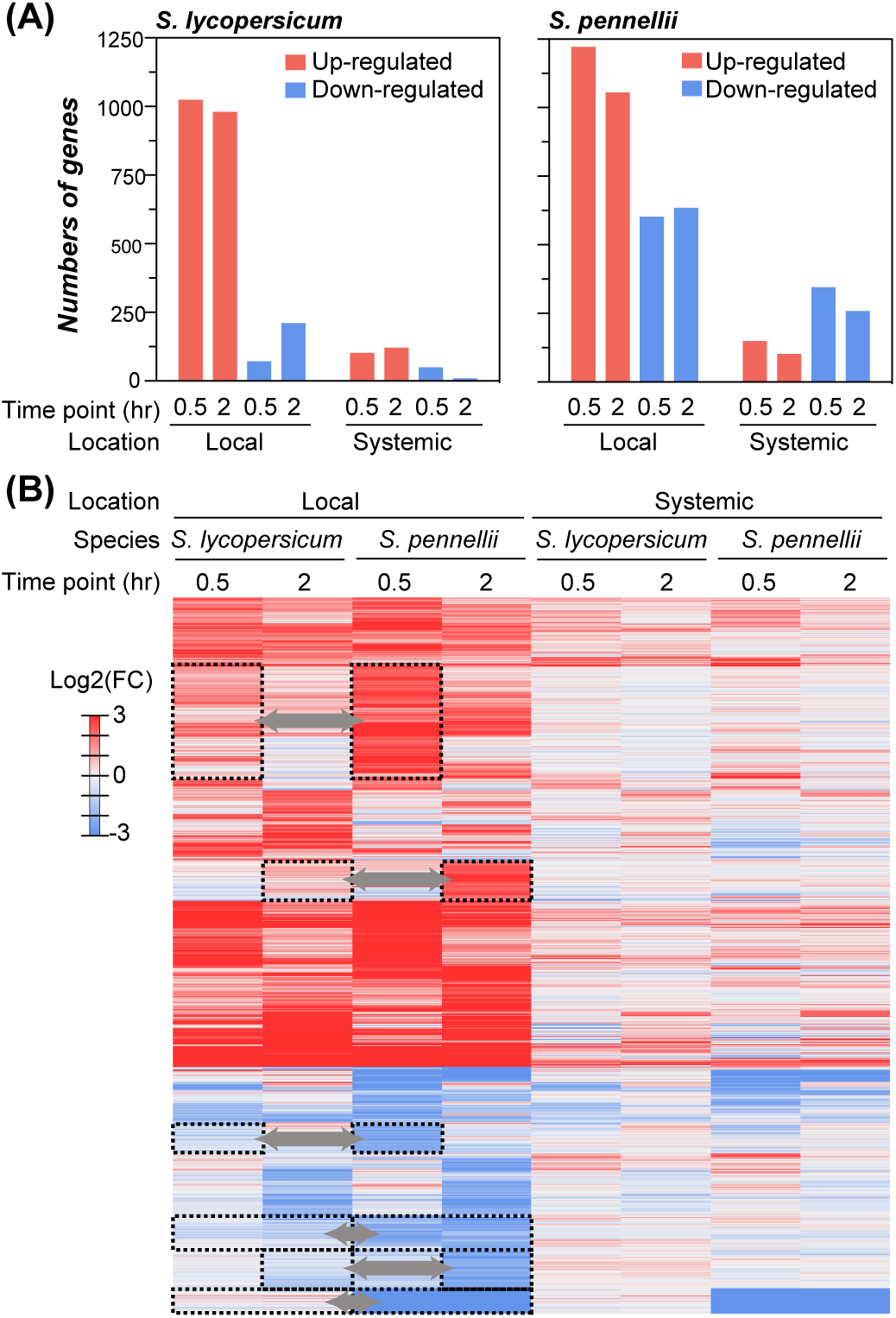
Similarities and differences in wound-responsive gene expression between tomato species. (A) Number of significantly differentially regulated genes upon mechanical wounding in local and systemic leaves of *S. lycopersicum* and *S. pennellii* for the time points (hr: hour(s)) post wounding indicated. (B) Bi-clustering of differential gene expression values (fold change, FC) of orthologous genes (rows) and location/species/time points (columns). Only orthologous genes significantly up- or down-regulated in ≥1 sample were included (n=2,183). Dashed boxes and arrows: indicating clusters of orthologous genes with inconsistent regulatory patterns across species in local tissues.

To assess in more detail how *S. lycopersicum* and *S. pennellii* differ in their wound response, orthologous genes that are wound-responsive (n=1,941) in any time point or tissue (i.e., local or systemic) in at least one species were compared. Hierarchical clustering of the overall expression patterns showed that the samples were clustered first based on the treatment location (local or systemic) and then by species (**Fig. 1B**), indicating that the spatial treatment has higher impact over the species origins or the duration of treatment on wound responsive gene expression. Although the overall patterns of up and down-regulation are similar between species, there are some important differences. In the local leaves at both time points, *S. pennellii* genes have higher amplitude of expression (higher absolute FC values) compared to *S. lycopersicum* orthologs (dashed boxes, **Fig. 1B**). Thus, *S. pennellii* apparently responds to wounding earlier and stronger than *S. lycopersicum*, which is consistent with the heightened tolerance to drought and salt in *S. pennellii* compared to *S. lycopersicum* [18, 24, 35, 36].

### Co-expression clustering and functions of wound-responsive genes

The overall transcript profile showed that wound-responsive genes differed significantly between species and could be classified into categories according to the time of treatment and spatial location of the response (**Fig. 1**). To further investigate how the wound response may have functionally diverged between species, we first categorized a wound-responsive gene from a species into one of 81 clusters representing different expression patterns, including whether the gene is up-regulated (“U”, red, log_2_(FC)>2), non-regulated (“N”, gray, |log_2_(FC)|<2) and down-regulated (“D”, blue, log_2_(FC)<-2) in response to wounding at a given time/location (**Fig. 2A**). These clusters are referred to as wound response clusters. For example, a gene is categorized in the UUDN cluster if it is up-regulated at both 0.5 hr and 2 hr time points in the local wounded leaf, down-regulated at the 0.5 hr time point in the systemic undamaged leaf, and not changed significantly in the 2 hr systemic response. Note that only a subset of clusters are shown in **Fig. 2A**; the remaining omitted clusters comprise <2% of wound-responsive genes (Supplemental Table S1) and are not discussed further. Among the seven major wound-induced clusters (red, **Fig. 2A**), the UNNN, NUNN, and UUNN clusters are the largest with >300 genes represented in both species (**Fig. 2B, Supplemental Table S1**). The total number of up-regulated genes in these three major clusters tended to be greater in *S. pennellii* than *S. lycopersicum*. Similarly, it was also apparent that the number of genes comprising the five major wound-repressed clusters (blue, **Fig. 2A**) was greater in *S. pennellii* (**Fig. 2B**). These general patterns of transcript abundance suggest that *S. pennellii* has more dynamic wound response, particularly in the case of down-regulated genes.

**Figure 2.**
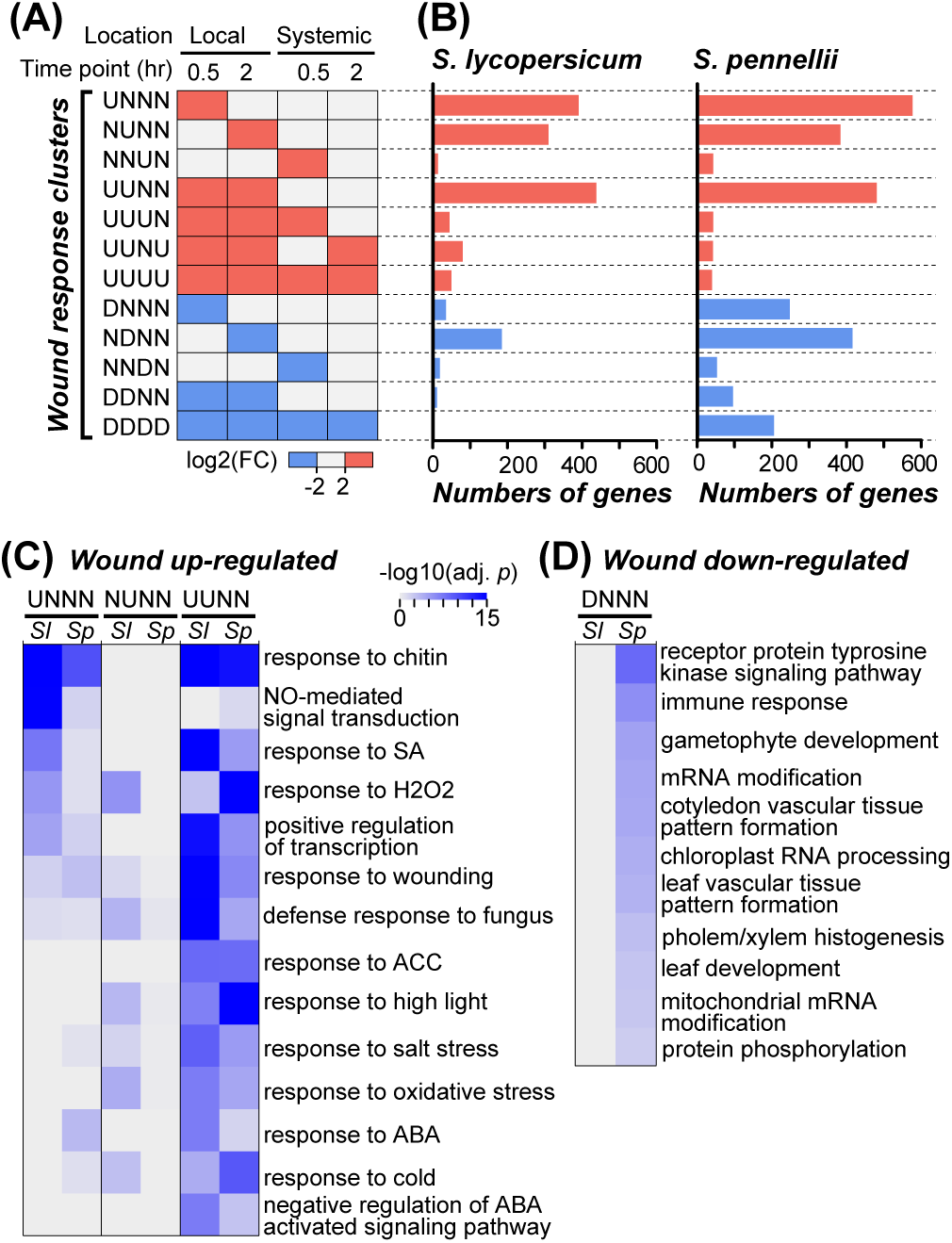
Number of genes and functional category enrichments in wound response clusters. (A) Definitions of wound response clusters. U (red): up-regulation. N (white): no significant change. D (blue): down-regulation. Only clusters with >40 genes in ≥1 species were shown. (B) Numbers of wound-responsive genes in the clusters shown in (A) for *S. lycopersicum* (left) and *S. pennellii* (right). Red: up-regulated clusters. Blue: down-regulated clusters. (C) Gene ontology biological process categories significantly enriched (false discovery rate adjusted p<1e-3) in *S. lycopersicum* (Sl) or *S. pennellii* (*Sp*) genes from wound up-regulated clusters. Deeper shades of blue indicate higher -log_10_(adjusted *p* value). (D) Categories enriched in genes from wound down-regulated clusters (adjusted *p*<1e-2).

Considering the differences in wound-responsive gene expression between *S. lycopersicum* and *S. pennellii* (**Fig. 1,2**), we assessed the function of wound-responsive genes in each wound response cluster with Gene Ontology (GO) and metabolic pathway annotations (see Materials & Methods). Wounding activates broad-spectrum defense responses in tomato [28, 29, 37]. We thus expected defense genes to be commonly up-regulated in these two species. Consistent with previous findings [28, 32], the wound up-regulated genes in local leaves, especially those in the UNNN and UUNN clusters, are significantly enriched in genes responsive to multiple biotic and abiotic stresses, including those mediated by the stress hormones salicylic acid (SA), abscisic acid (ABA), and ethylene (**Fig. 2C**). Notably, most biological processes are more significantly enriched in *S. lycopersicum* than in *S. pennellii* (**Fig. 2C**), regardless of time point in the local response (Supplemental Fig. S1). This result suggests that, although there are more wound-induced genes in *S. pennellii* (**Fig. 1**), wound stress induces a proportionally higher number of defense-related genes in the domesticated tomato than that in the wild species.

Although there is a large number of wound down-regulated genes (**Fig. 1**), only one cluster (DNNN) containing *S. pennellii* genes is significantly enriched in plant growth-related GO categories, including leaf development and vascular tissue formation (**Fig. 2D**). This is consistent with previous studies showing that wound-induced defense hormones, JA and SA, inhibit leaf development [38, 39] and the presence of tradeoffs between growth and stress tolerance in wild species [40]. We also found that genes in the UUNN and NDNN clusters in *S. pennellii* are significantly enriched in indole-3-acetic acid (IAA) degradation pathway and phylloquinone biosynthesis, respectively (Supplemental Fig. S2A,B). IAA is a growth hormone regulating processes that are impacted by defense responses [41], whereas phylloquinone is an integral part of the photosynthetic electron transport chain [42]. Changes in the expression of genes associated with enhanced IAA degradation and reduced photosynthetic efficiency are consistent with the antagonistic relationship between defense response and plant growth in *S. pennellii* (**Fig. 2D**).

Taken together, our findings show that wound-response genes can be categorized into a few dominant clusters (**Fig. 2A,B**). Because some orthologs have differing response to wounding (**Fig. 1B**), the identity and the enrichment test statistics of some GO categories and metabolic pathways also differ (**Fig. 2C,D**). Nonetheless, the number of GO categories and metabolic pathways enriched in genes up or down-regulated in either species was small. This is particularly true for *S. pennellii* down-regulated genes. Since only the orthologs were included in the gene set enrichment analyses (see Materials & Methods), the small numbers of GO categories recovered may be due to the lower gene number in a given cluster, which consequently decreases statistical power.

### Divergence of wound responses among orthologous genes

Previous work in maize and tomato has suggested that the domestication process or the adaptation to extreme environments may result in extensive changes in the transcriptional regulation of genes controlling relevant morphological and physiological traits [17, 18]. Our findings show that there are substantial differences in the wound-responsive expression of *S. lycopersicum* and *S. pennellii* genes, as well as differences in the biological processes represented by these genes (**Fig. 2, Supplemental Fig. S2**). One immediate question is to what extent the orthologous genes in these two species differ in their wound response. To address this question, we first assessed which putative orthologous genes (see Materials & Methods) have consistent wound response patterns (i.e. both orthologs are in the same wound response cluster) (**Fig. 3A**). These genes are referred to as consistent genes. Interestingly, relatively few orthologs belong to the same cluster (gray, **Fig. 3A**). Depending on the cluster, only 0-24% orthologs are considered consistent (**Fig. 3B**). These results show that 76-100% of the wound-responsive orthologous genes are in different clusters and thus differentially regulated between species. Note that this observation is not due to the small sample size of the clusters. Among the clusters with gene numbers >200 (UNNN, NUNN, UUNN and NDNN, **Fig. 3A**), >76% orthologous gene pairs belong to different wound response clusters (**Fig. 3B**). In assessing the orthologous gene expression patterns side-by-side between species, it is clear some orthologous pairs have substantially different responses (cyan and orange bar, **Fig. 3C-F**). For example, in the UNNN cluster (**Fig. 3C**), in 42% cases the *S. pennellii* orthologous genes are either in the NNNN cluster (dotted rectangle *a*, **Fig. 3C**) or in the UUNN cluster (dotted rectangle b, **Fig. 3C**). This suggests that wound responses have diverged among the majority of orthologs in the past ∼7 million years [43, 44].

**Figure 3.**
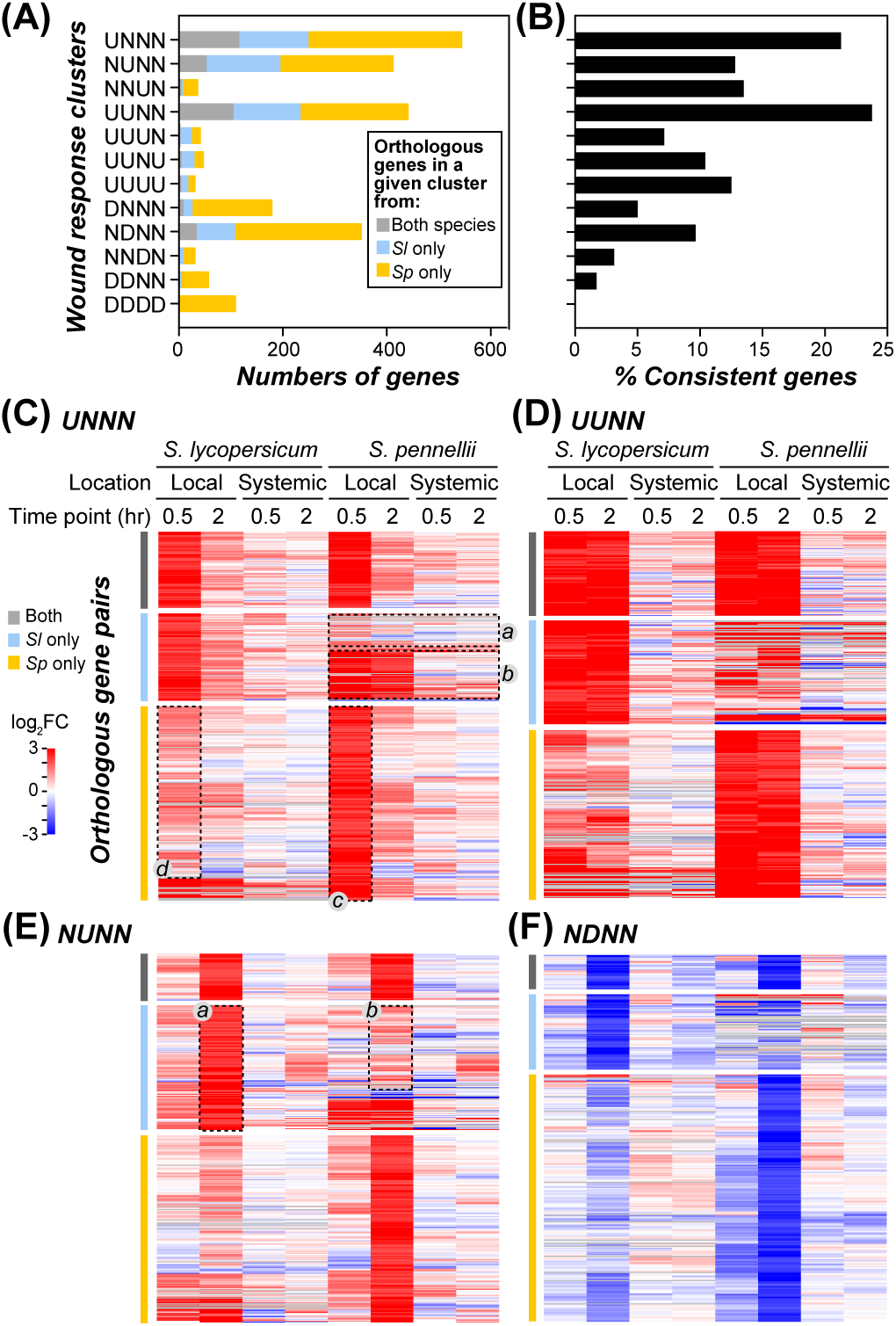
Divergence of wound responses among orthologous genes. (A) Number of orthologous genes in the wound response clusters as defined in **Fig. 2A**. Gray: orthologous genes from both species were in the same cluster. Cyan: the *S. lycopersicum* (*Sl*) ortholog is in the indicated cluster but not the *S. pennellii* one. Orange: the *S. pennellii* (*Sp*) ortholog is in the indicated cluster but not the *S. lycopersicum* one. (B) Percentage of the orthologous genes that are considered to have consistent regulatory patterns (in the same cluster) in each cluster. (C,D,E,F) Heatmaps showing the differential expression levels (log_2_(FC), FC: fold change) of orthologous genes in UNNN (C), UUNN (D), NUNN (E) and NDNN (F) clusters. The bars on the left of each heatmap are colored the same way as in (A). The dotted rectangles highlight differential expression patterns discussed in the main text.

To assess the extent to which the wound response differs between orthologous genes, we compared the wound-induced gene expression levels over the tested durations/tissues in the four largest clusters. In most cases, although they belonged to different clusters, both orthologs were responsive but at different levels. For example, for the orthologous gene pairs in UNNN cluster in which only the *S. pennellii* genes were significantly up-regulated (above threshold) at 0.5 hr in the local leaves (dotted rectangle c, **Fig. 3C**), the corresponding *S. lycopersicum* orthologs were also up-regulated but at levels below the threshold (dotted rectangle d, **Fig. 3C**). Similarly, in the NUNN cluster where only the *S. lycopersicum* orthologs were significantly up-regulated (dotted rectangle a**, Fig. 3E**), the expression of most corresponding *S. pennellii* orthologs was also induced but at levels below threshold (dotted rectangle *b*, **Fig. 3E**). This pattern is also true for down-regulated genes (cyan and orange bars, **Fig. 3F**). Given that most orthologs were wound-responsive but expressed at different levels, these results suggest that the ancestral genes of these orthologs were likely wound-responsive. Thus when the wound response of orthologous genes diverges, the divergence does not reflect complete loss or gain of response but more likely varying levels of responsiveness.

To this point our analysis focused on differential expression by comparing wounded leaves to unwounded, control leaves. Although induced gene expression is important for kickstarting defense systems in unfavorable environments [28, 29, 37], constitutive defenses also contribute to plant resilience to environmental stress [45]. Using the *S. lycopersicum* gene expression level as a reference, we identified 529 and 297 *S. pennellii* orthologs that are expressed to higher and lower levels, respectively, than cultivated tomato (Supplemental Fig. S3A). This finding indicates that significant differences in gene expression exist between the two species prior to wounding. For example, cuticular wax and cutin biosynthesis genes, CER6, CER8, MYB41 and SICUS2 [46-48], were expressed at higher levels in *S. pennellii* (**Supplemental Fig. S3B**), consistent with findings of earlier studies [24].

Given the expression levels are already different between the control samples, it is possible that a gene contributing to constitutive defense will have a consistently high expression level before and after wounding. To assess this, we also compared the gene expressions in wound-treated samples in both species against the *S. lycopersicum* unwounded control. A surprising pattern is that, if a *S. pennellii* gene has a significantly different expression level in unwounded control compared to that of the *S. lycopersicum* control, the *S. pennellii* gene in question will remain significantly different after wounding in both time points and in both local and systemic tissues (Supplemental Fig. S3A). This is consistent with the hypothesis that the basal level of defense response is stronger in *S. pennellii* [18, 24].

### Putative *cis*-regulatory sequences controlling wound-responsive gene regulation

The expression patterns in control and wounded tissue between *S. lycopersicum* and *S. pennellii* orthologous genes have diverged substantially, suggesting divergence of **cis*-* and/or *trans*-regulatory mechanisms central to controlling wound-responsive gene expression. At the *cis*-regulatory level, substitutions in CRE sites would alter TF binding affinity and thus contribute to expression differences [7, 21, 49, 50]. In tomato, studies of wound and JA-responsive gene expression have revealed that some promoter fragments and G-box motif regulate wound and JA responses [34, 51]. However, our knowledge of *cis*-regulation of wound-responsive expression is limited and, for the known CREs such as G-box, they have not been evaluated to globally assess their contribution to regulatory divergence across species. Thus to better understand the contribution of *cis* and *trans* regulatory divergence between these two species, we identified globally the CREs likely controlling wound-responsive gene expression for cross-species comparison with an enriched *k*-mer approach (an oligomer with the length *k* ≥ 5 base-pairs, see **Materials & Methods**).

Since CREs may be located in both the promoter and 5’UTR regions of a gene [32, 37], we queried whether a k-mer sequence is enriched near the transcriptional start sites (TSS, see **Materials & Methods**) of member genes in each cluster. Zero to hundreds of k-mers were found to have significantly enriched numbers of sites among clusters (**Fig. 4A**). These enriched k-mers are referred to as putative CREs (pCREs). The pCREs identified include ones that resemble known CREs relevant to the wound response, including ABRE, W-box and G-box [34, 52-55], which are discussed in later sections. To further assess how well these pCREs can jointly explain the wound response in each cluster, we applied a machine-learning algorithm, Support Vector Machine (SVM, see **Materials & Methods**), to predict wound-responsive expression of genes in a wound response cluster based on identified pCREs. Among the nine clusters with pCREs in *S. lycopersicum* and/or *S. pennellii* (**Fig. 4A**), the wound-response prediction models based on pCREs performed significantly better than randomly expected (boxplots vs. gray spot, Wilcoxon signed rank test, all p<0.027; **Fig. 4B**). These results showed that our approach could efficiently identify short sequences resembling CREs because they are predictive of wound response in multiple clusters. In addition, the pCREs from clusters involving wound-induced expression (e.g. UNNN (red) and UUNN (orange), **Fig. 4C**) tend to be located within 500 bp upstream of the TSS, consistent with the finding that plant TFs tend to bind preferentially in the upstream region close to TSSs [56, 57].

**Figure 4.**
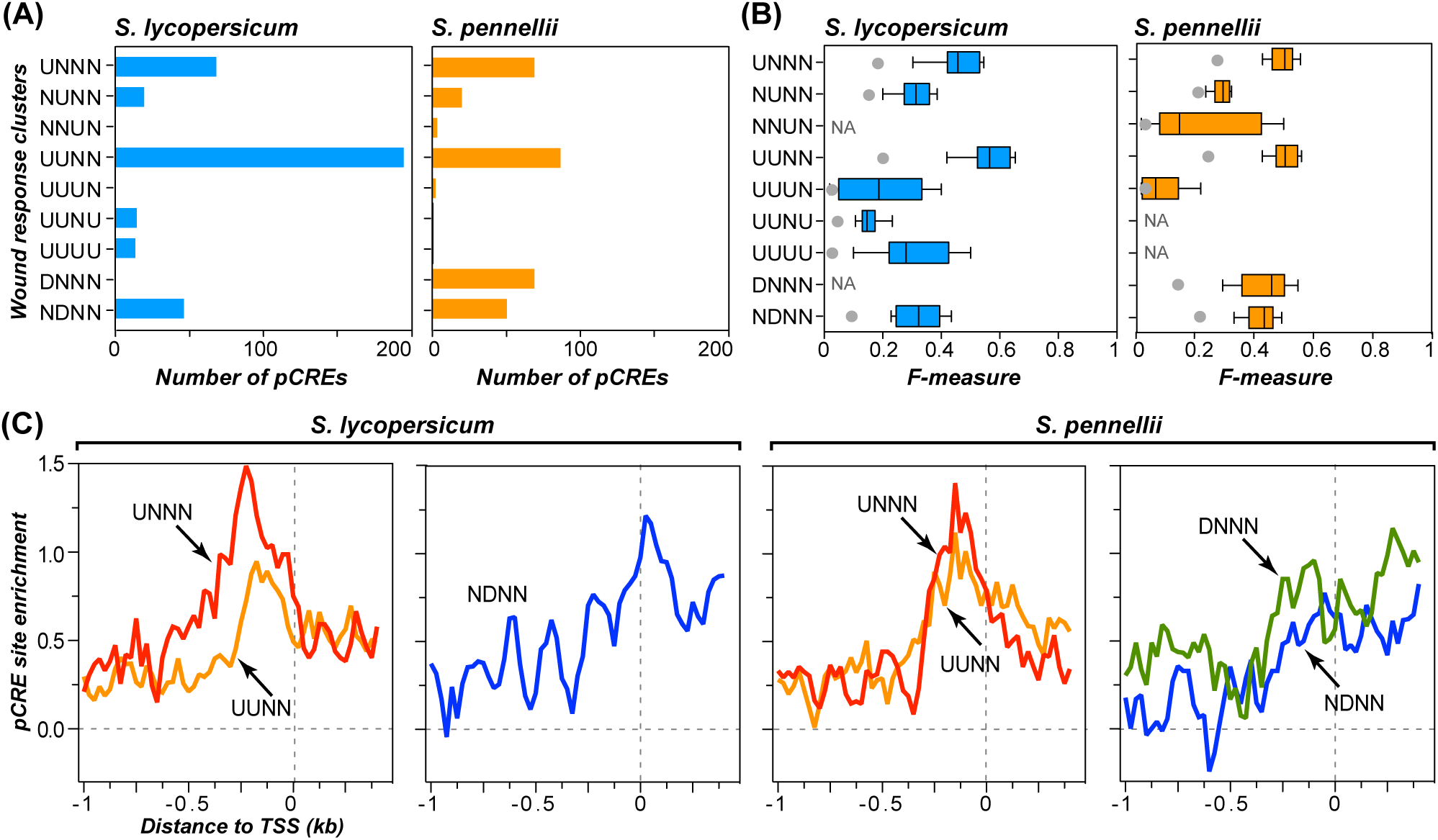
Evidence indicating biological relevance of putative CREs. (A) Number of putative CREs (pCREs) identified through the *k*-mer pipeline (see **Materials & Methods**) for each wound response cluster in *S. lycopersicum* (blue) or *S. pennellii* (orange). Only the clusters with pCREs in ≥1 species are shown. (B) Boxplot showing the wound response prediction performance (F-measure) based on a model using pCREs identified from genes in a wound response cluster. F-measure: the harmonic mean of precision (proportion predicted correctly) and recall (proportion true positives predicted). The maximum F-measure is 1, indicating a perfect model. For each wound response cluster, 10 F-measures were calculated from 10-fold cross validation and shown as a boxplot. Gray dot: the average F-measure of 10,000 random predictions indicating the performance of a meaningless model. NA: not applicable since no pCRE was found in the cluster. (C) Enrichment of sites of pCREs identified from four different clusters. For each pCRE, the degree of enrichment of its sites around transcription start sites (TSSs) was represented as the log_2_ ratio between pCRE site frequencies of genes in a cluster and frequencies of the same pCREs in genes not responsive to wound. This log ratio was generated for each pCRE in the region from 1 kb upstream to 0.5 kb downstream of transcription start sites (TSSs) with a sliding window of 100 bp and a step size of 25 bp. For each cluster, the median log ratios of all pCREs identified from the cluster in question was shown.

In contrast to pCREs involved in up-regulation, the pCREs identified in wound down-regulated clusters (NDNN (blue) and DNNN (green), **Fig. 4C**) tend to be located around TSSs, including 5’UTRs. This is similar to the 5’UTR of excision repair cross complementation group-1 gene in human that contains binding sites for a transcription repressor [58]. Similarly, the cyclin D1 Inhibitory Element within the 5’UTRs represses the expression of the human cyclic D1 gene in an-age-dependent manner [59]. Nonetheless, we cannot find pCREs in most clusters relevant to down regulation (DDDD, DDNN, NNDN), suggesting an important role for post-transcriptional regulation such as transcript turnover [60] in control of these genes. Taken together, the pCREs identified are predictive of wound-responsive gene expression in most clusters and have a position bias resembling the known TF binding sites, suggesting that they are authentic *cis*-elements in regulating gene expression. In the next sections, we compared how these pCREs and their locations had diverged between species and how such divergence was relevant to the divergence in wound response.

### Conservation of putative CREs between tomato species

To determine if pCREs are conserved between *S. lycopersicum* and *S. pennellii*, we first asked: if a pCRE is enriched in a wound response cluster in one species, is the same pCRE enriched in the same cluster in the second species? If a pCRE is consistently enriched in the same cluster across species, it would be an indication that the pCRE in question was a component of a conserved wound response regulatory program. We found that 35-66% of pCREs in UNNN, UUNN and NDNN clusters were common between species. For example, among pCREs identified from *S. pennellii*, 37 (52.8%), 58 (65.9%), and 20 (39.2%) were also enriched in the *S. lycopersicum* UNNN, UUNN and NDNN clusters, respectively (**Supplemental Table S2**). Although the wound responses have diverged significantly, our naïve expectation was that the underlying regulatory programs would be conserved. The relatively low degree of conservation was unexpected and could be because there were distinct TFs between species that binds different sets of *cis*-regulatory sites controlling wound response. Alternatively, this pattern may reflect the degeneracy of TF binding sites [61]. For example, TF-regulated genes within a cluster remained the same (orthologous) between species but these TFs bind *cis*-regulatory sites with subtle differences [61].

To assess the above possibilities, we first defined species-specific pCREs as those that were enriched only in *S. lycopersicum* or *S. pennellii* genes within a given cluster, and then asked whether these two sets of species-specific pCREs could be bound by TFs from the same family. The rationale is that, if the species-specific pCREs for a wound response cluster are not bound by TFs of the same family, then they must have diverged *trans* regulatory mechanisms. Using *in vitro* TF binding data [50], a threshold distance was defined where TFs from the same family had distances above threshold (see **Materials & Methods**) and used to define pCRE sub-groups (**Fig. 5A** and **Supplemental Fig. S4-S5**). In each sub-group, the pCREs were likely bound by TFs of the same family. Therefore, if a sub-group contains pCREs from both species, the pCREs within this sub-group were defined as “conserved” across species. In contrast, if all pCREs in a sub-group comes from only one species, these pCREs are regarded as “non-conserved” (asterisk-labeled, **Fig. 5A**). For example, among pCREs that are enriched in UNNN wound responsive genes from at least one species, they can be divided into 31 sub-groups (**Fig. 5A**). Using the UNNN, UUNN, and NDNN clusters that are among the largest as examples, 84-100% of the pCREs were in sub-groups containing pCREs from both species (**Fig. 5A, Supplemental Fig. S4-S5, Supplemental Table S2**).

**Figure 5.**
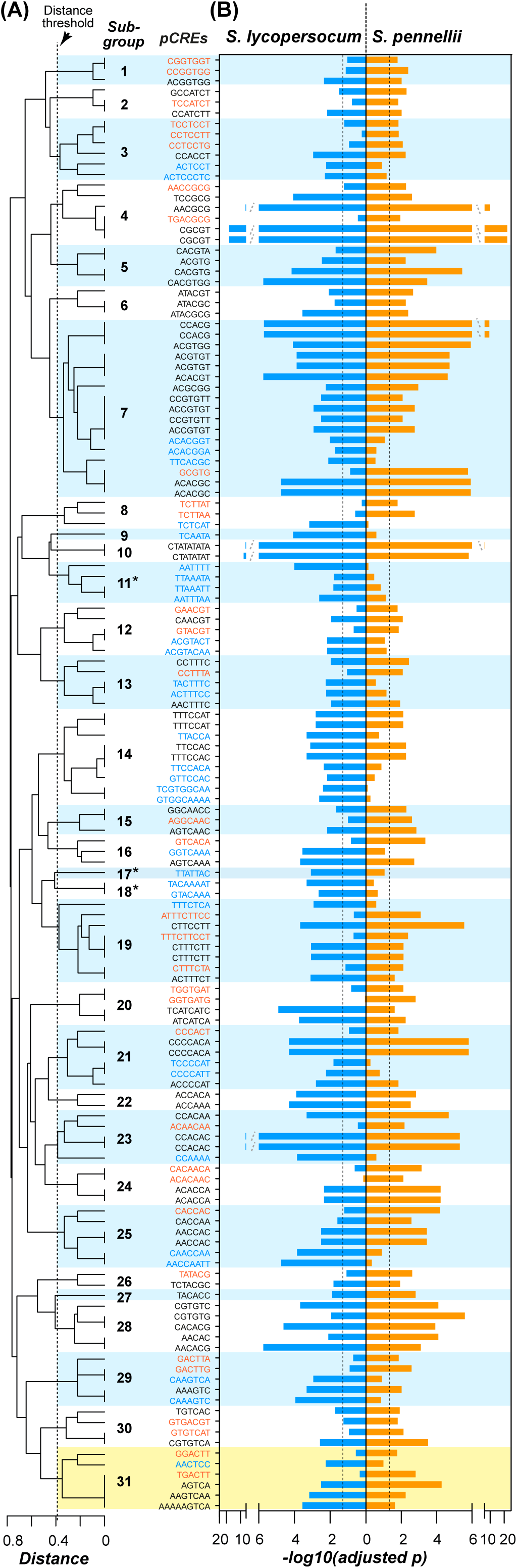
Differential enrichment of pCREs in UNNN cluster genes from two tomato species. (A) Dendrogram showing the distances between the pCREs identified from UNNN cluster genes from in *S. lycopersicum* only (blue), *S. pennellii* only (orange), or both species (black). The dotted line indicates the threshold distance defined based on the 95^th^ percentile distances between binding motifs of TFs from distinct families and defines multiple pCRE sub-groups (numbered) where each sub-group contains pCREs likely bound by TFs of the same family (distance threshold=0.39). Subgroups with pCRE from only one species are labeled with asterisks. (B) Degrees of pCRE site enrichment in *S. lycopersicum* (blue) and *S. pennellii* (orange) UNNN genes. Adjusted p-value: multiple-testing corrected p-value. Dashed line, adjusted *p* <0.05. Yellow box: pCREs similar to the W-box element.

Taken together, 35-66% pCREs were enriched in the same wound response clusters across-species. Thus >34% pCREs could be defined as species-specific due to this differential enrichment. In addition, >74% (depending on the wound response cluster) of these species-specific pCREs were conserved between *S. lycopersicum* and *S. pennellii* in the sense that they could be bound by TFs of the same family (Supplemental Table S2). Therefore, a substantial number of TFs from the same family were important for regulating wound response in both species. We note that <26% of the species-specific pCREs were differentially enriched and sufficiently distinct that they may be bound by unrelated TFs, suggesting *trans* regulatory divergence (**Fig. 5A**, **Supplemental Fig. S4-S5, Supplemental Table S2**). Nonetheless, the sub-groups were defined to ensure pCREs bound by TFs from the same family can be correctly identified. Thus the 26% represents the upper bound in terms of the degree of *trans* regulatory divergence in wound response between these two species.

To further investigate which TFs may bind to the pCREs in different sub-groups (Supplemental Fig. S6A-C), the pCREs in a sub-group were merged and compared to 279 CREs known to bound by 256 plant TFs [50] (see **Materials & Methods**). Among 130 conserved pCREs in the UNNN cluster, some were similar to the known stress-responsive CREs (“Conserved”, up panel, **Supplemental Fig. S6A**). For example, the CACGT[A/G] representing the merged pCREs in a given subgroup is similar to the G-box elements bound by basic leucine-zipper (bZIP) and basic helix-loop-helix (bHLH) transcription factors (upper panel, **Supplemental Fig. S6A**), which are known to be important for wound- and JA-responsive gene expression [55, 62, 63]. The TGACTT pCRE is similar to the W-box bound by WRKY factors (upper panel, Supplemental Fig. S6A) that also regulates wound and JA responsive gene expression [64]. In UUNN clusters, the pCREs identified were similar to the stress-responsive CREs. The consensus sequences (ACGTGA, G[A/T]ACGT, TTGAC[C/T]) are similar to the known defense-responsive elements, G-box, ABRE and W-box and, that are bound by bHLHs, WRKYs and bZIPs (**Supplemental Fig. S6B**) [34, 52-55]. AATTTA is similar to the binding motif of AT hook factors, which regulate plant immune and JA response (**Supplemental Fig. S6B**) [65, 66]. These results are consistent with our findings that wound treatment triggered defense response against multiple stresses (**Fig. 2C**), and further support the relevance of these pCREs.

### Relationship between pCRE conservation and gene regulation across species

In the previous section, we observed that pCREs identified showed different properties when analyzing the conservation (i.e. recognized by the same TF family) and enrichment in genes between species (**Fig. 5 and Supplemental Fig. S4-S5**). One immediate question is if the conserved or the species-specific pCREs contribute more significantly to wound transcriptional response. To address this, we classified the pCREs into 3 types: Type I (pCREs conserved and enriched in both species), Type II (pCREs conserved across species but enriched in a species–specific manner) and Type III (pCREs non-conserved and enriched in a species–specific manner) (**Supplemental Fig. S6D**). These three types of pCREs were then used to build machine learning models (see **Materials & Methods**) for predicting wound-responsive expression of genes in a wound response cluster. We found that models built with Type I pCREs were in most cases the best at predicting wound response in both species (gray box, **Supplemental Fig. S6E**), suggesting that these pCREs are components of conserved regulatory mechanisms across species. Type II pCREs predicted wound response well within species but not across species (cyan and pink, **Supplemental Fig. S6E**), supporting the observation of *trans*-regulatory divergence. We should note that, except the NDNN clusters in *S. pennellii*, the prediction performance of Type II and III pCREs were not as accurate as the Type I pCREs (**Supplemental Fig. S6E**). In addition, only some of the clusters could identify the Type III pCREs (**Supplemental Fig. S6E**). This suggests that the conserved *cis*-regulatory elements play a more central role in wound-responsive transcription in both tomato species, and that distinct pCREs may contribute to differential gene expression in each species.

### Distinct preference of pCREs between wild and domesticated species

We found that 84-100% pCREs could be defined as conserved across species based on the definition that pCREs bound by TFs from the same family and controlling wound response in a wound response cluster were found in both species (**Fig. 5A, Supplemental Table S2**). However, this definition of conservation underestimates the degrees of *trans* regulatory divergence because these pCREs were likely bound by different TFs of the same families. Consistent with this, only 35-66% of these conserved pCREs were significantly enriched in both species (**Fig. 5B, Supplemental Table S2**). This shows that a substantial number of pCREs were enriched in a species-specific manner. As mentioned in the previous section, an explanation is that, although these conserved pCREs are highly similar (belong to the same sub-group, **Fig. 5**), the regulatory factors in these two species may prefer similar but distinct pCRE sequences [11, 67]. To test this hypothesis, we asked if pCREs that belong to the same sub-group (targeted by TFs of the same family) have different degrees of enrichment in *S. lycopersicum* and *S. pennellii*. Consistent with our explanation, pCREs are specifically enriched (at the 5% significance level) in only one species in 21 of 28, 39 of 50, and 19 of 20 sub-groups in UNNN, UUNN and NDNN clusters, respectively (**Fig. 5**, **Supplemental Fig. S4-S5**).

For example, in the UNNN cluster the consensus sequence of six pCREs in the 31^st^ sub-group is TGACTT (yellow box, **Fig. 5**) that is similar to the W-box (TTGAC[C/T]) recognized by WRYK TFs that mediate biotic and abiotic stress responses [58]. Among these six pCREs, TGACTT and GGACTT were enriched specifically in *S. pennellii* whereas AACTCC was enriched in *S. lycopersicum*. In addition, we found that the “preferred” pCREs (i.e. the pCRE with the most significant degrees of enrichment in each species in each subgroup) are frequently distinct (**Fig. 5B**). For example, in this 31^st^ sub-group AGTCA (reverse complement, TGACT) was the most significantly enriched in *S. pennellii*, whereas AAAAAGTCA (reverse complement, TGACTTTTT) was the most significantly enriched in *S. lycopersicum* (**Fig. 5B**). Similar phenomena were also observed for the pCREs identified in the UUNN cluster (Supplemental Fig. S4). These results showed that, although these pCREs are similar to each other and potentially bound by TFs of the same family, they were differentially enriched between species, possibly due to diverged binding preference of TFs between *S. lycopersicum* and *S. pennellii*. This is in line with the previous study that the orthologous LEAFYs from algae and land plants as well as the orthologous TFs from fruit fly and human showed differences in binding site specificity due to subtle amino acid differences [11, 68].

Taken together, although *S. lycopersicum* and *S. pennellii* may employ conserved pCREs and similar sets of TFs to control wound-induced gene expression, there are distinct preferences for pCRE sequences that reflect *trans* regulatory divergence (**Fig. 5**, **Supplemental Fig. S4-S5**).

### Turnover of putative CRE sites between orthologous genes and their association with gene regulation

Our findings so far indicated substantial conservation of *cis*-regulatory elements between domesticated and wild tomato species (**Fig. 5**, **Supplemental Fig. S4-S5**). In addition, we found extensive variation of wound-responsive gene expression among orthologous genes (**Fig. 3**). These differences may result from minor changes in CRE sequences, leading to differences in TF binding specificity (**Fig. 5**). Alternatively, the wound response divergence between orthologs may be the consequence of differential turnover (i.e. the gain and loss) of the *cis*-regulatory sites within orthologous regions [3, 7]. We next determined the extent to which these *cis*-regulatory sites were conserved or turned over across species and their association with gene expression divergence. Based on the relative position of the sites located in regulatory regions of orthologous gene pairs, the sites of a given pCRE were categorized into “shared”, “specific”, “compensatory” and “other” types (**Fig. 6A**; see **Materials & Methods**). Since the “compensatory” and “other” types accounted for small portions of the pCRE sites (**Supplemental Fig. S7**), we focused on the “shared” and “specific” pCRE types.

To summarize the degree of conservation of the sites of each pCRE identified from various wound response cluster (**Fig. 4A**), a conservation likelihood (*L_c_*) for each pCRE was computed by calculating the log2 ratio between the proportion of sites that are shared and the proportion of sites that are specific (see **Materials & Methods**). Thus, a higher *L_c_* indicated a higher degree of enrichment of shared sites relative to that of specific sites. A pCRE with a higher *L_c_* was considered more conserved than that with a lower *L_c_*. First, to assess if the conservation of pCRE sites was correlated with the consistency of the wound response between orthologs, we compared the *L_c_* values for the orthologs with consistent wound response and for those with inconsistent patterns. Using the UNNN cluster as an example (left panel, **Fig. 6B**), we found that the sites of pCREs in orthologous gene pairs with consistent wound response patterns (median *L_c_*=0.57) had significantly higher *L_c_* values than sites in orthologous pairs with inconsistent patterns (median L_c_=0.28, Mann-Whitney U test, p=2.1e-04). The same was true when comparing pCRE sites in genes with consistent patterns against sites found in the non-responsive orthologous genes (median *L_c_*=-0.58; p<2.2e-16) (left panel, **Fig. 6B**). Similar results were also observed for the pCREs in the UUNN cluster (middle panel, **Fig. 6B**). Taken together, these results imply that in UNNN and UUNN clusters, the orthologs with consistent gene regulation tend to have more conserved pCRE sites, indicating that, as expected, conservation of pCREs sites contribute to a conserved wound up-regulated response across species.

**Figure 6.**
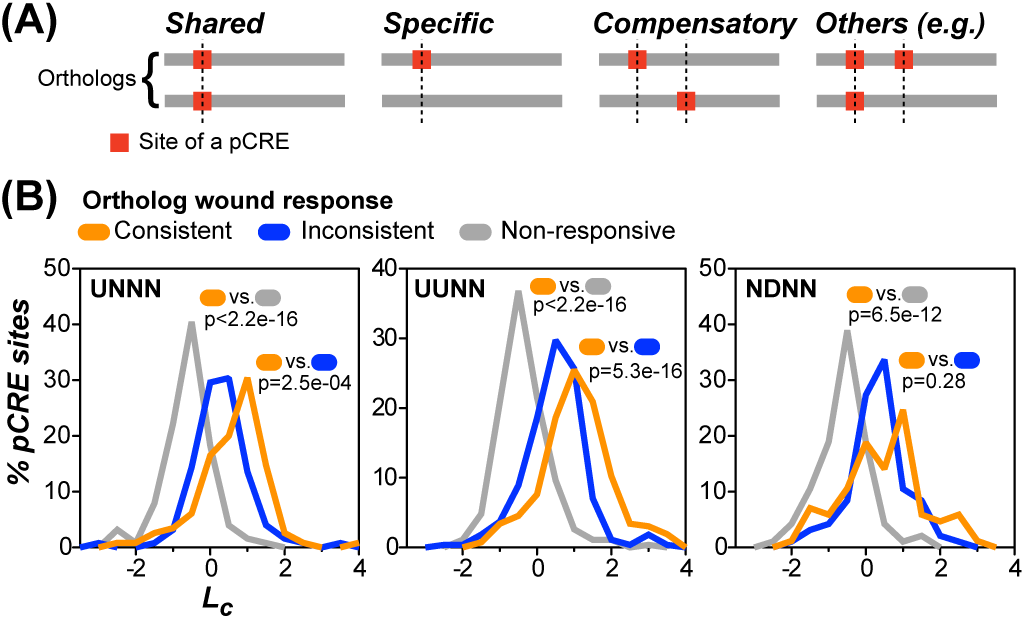
Relationships of pCRE site turnover and wound responsive gene expression between orthologs. (A) Types of pCRE sites. Shared: the sites of a pCRE are present in both orthologs and located at the same position. Specific: the site of a pCRE is present only in one ortholog but not the other. Compensatory: the sites are present in both species but in different locations. Others: any situation that does not belong to the previous three types. Gray line: the defined regulatory regions from the orthologous gene pairs (See **Materials & Methods**). (B) The conservation likelihood of a pCRE (*L_c_*) in the UNNN (lef panel), the UUNN (middle panel), and NDNN (right panel) clusters. For a pCRE, its *L_c_* is defined as the log2 ratio between the proportions of sites that shared and those that are specific (see **Materials & Methods**). The *L_c_* for each pCRE was evaluated using orthologous gene pairs with consistent (belong to the same wound response cluster, orange) and inconsistent (belong to different clusters, blue) wound responses, as well as orthologous genes that are not responsive to wounding (Non-responsive, gray). *P-*values: testing whether the likelihood scores generating based the blue or gray datasets differ from the orange one (Mann-Whitney U test).

In contrast, in wound down-regulated cluster (NDNN), although the pCRE sites in orthologous pairs with consistent wound response patterns have higher median *L_c_* value (0.46) than those with inconsistent patterns (0.36), the difference was not significant (p=0.28; right panel, **Fig. 6B**). Thus, for down regulation, the degree of pCRE site conservation was not generally correlated with the differences in wound transcriptional response across species. One possibility is that in addition to the conserved pCREs, the non-conserved pCREs between species may be also involved in differential gene expression. Consistent with this, the models for predicting gene membership in the NDNN cluster performed significantly better when non-conserved pCRE sets (Type III) were used compared to random predictions (light blue and light pink box vs. gray spots, **Supplemental Fig. S6E**). Another possibility is that, compared to environmentally induced expression, regulatory events beyond transcriptional initiation (e.g. transcript turnover rates) may play a more significant role in down regulation. This is consistent with our machine learning findings that using pCREs to predict genes with the NDNN expression pattern tends to have poor performance compared to UNNN and UUNN up-regulated clusters (**Fig. 4B**).

Taken together, these findings suggest a positive correlation between the degrees of pCRE site conservation and the conservation of wound up-regulated gene expression between wild and domesticated species. On the other hand, the lack of a significant correlation observed in wound down-regulated genes suggests, except the conserved pCREs, the pCREs distinct between *S. lycopersicum* and *S. pennellii*, or post-transcriptional mechanisms may be also involved in the differential transcriptional repression across species.

## CONCLUSION

In this study, we investigated the patterns and mechanisms of transcriptional divergence of environmental stress response in a wild and a domesticated tomato species. Specifically, our analyses focus on wound-responsive gene expressions and the *cis* and, indirectly, the *trans* regulatory components regulating wound response. Despite the recent (∼2.7 - 7 million years ago) evolutionary divergence between the wild *S. pennellii* and domesticated *S. lycopersicum* species [43, 44], we found that the wound-responsive expression patterns of the orthologous genes diverge due to the combined action of natural and artificial selection. In addition, we characterized the pCREs significantly associated with the wound regulation. pCREs identified in *S. lycopersicum* and *S. pennellii* were predictive of gene expression. Among these sequences, conserved pCREs could better explain gene regulation between species than those that are non-conserved. This is in line with previous studies that the TF-binding specificity evolves slowly and are highly conserved amongst Drosophila, mouse and human [10, 11]. Intriguingly, our studies found that the non-conserved pCREs and the pCREs distinctly preferred in one species partially explain wound response within species, indicating the divergence of *trans* regulatory components occurs after speciation.

Our finding of correlation between the turnover of the pCRE site and the expression divergence of orthologous genes, especially in wound up-regulated clusters, further support the evolutionary conservation of CREs for wound response in tomato. We should emphasize that, although the correlation is apparent, it is far from perfect. Specifically, some pCRE sites enriched among wound responsive genes displayed high degrees of conservation between orthologous pairs with inconsistent wound response patterns. One possibility is that these conserved pCREs in orthologs with inconsistent patterns are still regulating weaker wound response. This is because the wound response clusters were defined based on threshold differential expression. As discussed earlier, some orthologous genes were classified into different clusters despite similar but significantly weaker response (**Fig. 3**). We should also point out that the *L_c_* distribution of some orthologs with consistent wound response may also be low, indicating that consistent expression patterns cannot be easily attributed to the pCREs analyzed. This highlights the complexity of the transcriptional regulatory systems and the need for studies to further ascertain the mechanistic basis of stress response conservation and divergence. Lastly, among these sites located in the regulatory regions, it is possible that only part of them are the *in vivo cis*-regulatory sites which would be determined by the environmental factors such as the chromatin structure or the GC-content on the surrounding regions [69, 70]. Thus, studies aimed at characterizing the chromatin structure and identifying the combinatorial relationship between CREs will help to reveal the evolution of the *cis*-regulatory codes.

In conclusion, our study provides global comparative analyses to connect the divergence of pCREs and turnover of *cis*-regulatory sites to gene expression divergence between species and orthologous genes. The comparison of pCREs predictive of the wound response revealed the presence of both *trans*-regulatory conservation and divergence. The correlation between the turnover of the *cis*-regulatory sites and the differential expressions of orthologs discovered the *cis*-regulatory divergence underlying gene expression variation. Collectively, these findings advance our understanding of the mechanistic basis underlying the stress-responsive gene expression divergence across species.

## METHODS

### Plant materials and growth conditions

*S.lycopersicum* cv Castlemart was used as the domesticated species. Seed for the wild species, *S. pennellii* (LA0716), was obtained from Tomato Genetic Resource Center (UC Davis) and grown on Jiffy-7 peat pot (Hummert International, Earth City, MO) in a growth chamber under 16 h light (6:00 to 22:00, 200 μmol m^-2^ s^-1^)/ 8 h dark cycle at 28°C. Three to four week-old plants with 3 to 4 expanded true leaves were used for wound treatment as previously described [71]. For wound elicitation, the lower (older) two leaves were crushed with a hemostat across the midrib of all leaflets. All wounding was performed in the morning (8:00-9:00), 2 hr after the start of the light cycle. At the indicated time points, leaflets were excised with a razor blade and immediately frozen in liquid nitrogen. Damaged leaflets (local, older leaves) from the first and second leaves, and undamaged leaflets (systemic, younger leaves) from the third and fourth leaves were collected separately. Control leaves were harvested from a set of unwounded plants grown side-by-side with the set of wounded plants. Three biological replicates (i.e., three separate sets of plants) were harvested for each treatment and time point. Total RNA was isolated from frozen leaf tissue using a RNeasy Plant Mini Kit (Qiagen, Germantown, MD, USA). RNA sequencing (100-bp paired-end reads) was performed with the Illumina HiSeq2500 platform in the Michigan State University Research Technology Support Facility.

### Sequencing data processing

To map the RNA-seq reads and determine the gene expression level, the reference genome sequences and gene annotation of *Solanum lycopersicum* (ITAG2.4) and *S. pennellii* (Spenn_v2.0) were retrieved from Sol Genomics Network (https://solgenomics.net). The *S. pennellii* gene annotation was further re-annotated through Maker-P module [72]. The cumulative distribution plot of AED (Annotation Edit Distance), which provides the measure of how well the annotations are supported by the EST, protein and RNA-Seq evidence [72], showed that the MAKER-mediated version performed better compared to the Spenn_v2.0 version (red vs. black lines, **Supplemental Fig. S8**). To further evaluate MAKER-P performance, we focused on genes annotated in both datasets (n=22,292) (Supplemental Table S3). Among the genes with one-to-one relationship (i.e. overlapped genic regions) between Spenn_v2.0 and MAKER-P annotated version, the gene models annotated by MAKER-P had higher AED values than in Spenn_v2.0 version (green vs. gray lines, **Supplemental Fig. S8**). These results showed that MAKER-P improved the gene annotations of *S. pennellii;* thus the MAKER-mediated version was adopted in this study.

The paired-end RNA reads were trimmed with Trimmomatic (default setting except leading=20, trailing=20 and minlen=20) [73] and mapped to the genome with Tophat2 (version 2.0.8) [74]. Transcript levels of annotated genes were calculated with Cufflinks (version 2.1.1) [75] and shown as FPKM (Fragments Per Kilobase per Million fragments mapped). The numbers of raw, quality filtered and mapped reads and the sequencing coverage are reported in Supplemental Table S4.

To evaluate the reproducibility of gene expressions among replicates and the similarity of gene expression profiles among treatments, the Spearman’s rank correlation coefficient was determined by pairwise comparison of gene expression between 2 samples. The distance (1-Spearman’s rank correlation coefficient) was used to generate the dendrogram through hierarchical clustering function with “complete” method (**Supplemental Fig. S9**). Except the samples from the locally 2-hr wound-treated treatments in *S. pennellii* and from the systemically 0.5-hr wound-treated treatments in *S. lycopersicum* (bold red and bold black, **Supplemental Fig. S9**), the three replicates of a given treatment in one species were clustered together, showing the gene expression profiles were similar and reproducible among replicates (**Supplemental Fig. S9**). We noted that the replicate #2 and #3 of the locally 2-hr wound-treated samples in *S. pennellii* were clustered with samples from *S. lycopersicum* (red, bold, **Supplemental Fig. S9**) and had higher mapping percentages when reads were mapped to *S. lycopersicum* genome than *S. pennellii* one (**Supplemental Table S4**). These results suggest that the reads in these two *S. pennellii* samples contaminated with the reads from *S. lycopersicum*. Thus, only the replicate #1 of 2-hr wound-treated and local samples in *S. pennellii* was included and represented for the given condition. For the systemically 0.5-hr wound-treated sample in *S. lycopersicum*, the replicate #2 was clustered away from the other 2 replicates (black, bold, **Supplemental Fig. S9**), showing that lower expression correlation with the other 2 replicates in pair-wise comparison. Therefore, the replicate #2 was excluded and only the replicate #1 and #3 was included for the downstream analyses.

To identify the significantly wound-responsive genes, only protein-coding genes with the value of “FPKM_confidence_low” >0 were considered. The transcript abundance of control and wound-treated samples were compared with EdgeR [76]. Any genes with false discovery rate adjusted p<0.05 [77] and with 4-fold difference in RNA level between wound and control (unwounded) samples was considered to be wound-responsive and included for the analyses.

### Identification of putative *cis*-elements and prediction for wound response

Wound-responsive genes are categorized into the different regulatory clusters depending on the levels of differential gene expression in the indicated points as defined in **Fig. 2A**. Genes are regarded as nonresponsive genes if their log2 fold-change (FC) values in any wound-treatment samples, compared to the control one, are between 1.2 and 0.8 (n=3,619 in *S. lycopersicum*) and between 1.3-0.7 in *S. pennellii* (n=3,138). Noted that the FC range of defining the nonresponsive genes were arbitrarily determined so that the number of nonresponsive genes were similar between species. To identify the putative *cis*-regulatory elements (pCREs) associated with wound response (**Fig. 4A**), a *k*-mer (oligomer with the length of k) pipeline was established by examining the frequency enrichment of a *k*-mer sequence in the regulatory region among the genes of a given wound response cluster compared to the nonresponsive genes and determining the adjusted *p-values* through Fisher’s exact test and multiple testing (Benjamini-Hochberg method) [77]. Here, the regulatory region is defined as the region ranging from upstream 1 kb to downstream 0.5 kb of transcription start site (TSS).

Since the *cis*-regulatory element range from 5 nt to ∼30 nt [78], this *k-mer* pipeline includes several steps to discover the pCREs with various sequence lengths. Step 1: a set of all possible 5-mer oligomers was evaluated for the enrichment. Only the 5-mers with significant enrichment (adjusted *p-values*<*0*.05) were retained for the next step. Step 2: the sequence of the significantly enriched 5-mers from step 1 was extended with 1 nt of A, T, C or G in either direction, examined for their enrichment and retained if they are significantly enriched (adjusted *p-values*<*0.05*). The step was literally repeated until no extended *k*-mer sequence was found to be significantly enriched among the regulated genes. Noted that if two *k*-mers were both significantly enriched and one *k*-mer sequence exactly matches the other one, only the one with lower adjusted *p-value* was retained. Step 3: as described in step 1, but start with a set of all possible 6-mers. The significantly enriched 6-mers were combined with the set of the *k*-mers identified from step 2. Step 4: as described in step 2, but start with the set of *k*-mers from step 3. Finally, the set of *k*-mers significantly enriched in the indicated wound response cluster was determined and considered as pCREs (**Fig. 4A**).

The support vector machines (SVM) method that allows predicting of wound response of a gene based on a set of pCREs was performed using the LIBSVM implementation of the SVM method through the Weka wrapper with the parameters described previously [79]. The pCREs were used as attributes whereas the binary status of genes with/without wound regulation was the class we wanted to predict. For training the predictive models for each regulatory pattern, the genes of the given clusters are positive examples whereas the non-responsive genes are negative examples.

### Sequence similarity of putative CREs between species and to the known TF binding motifs

To identify the pCREs whose sequences are more significantly similar than expected between TF families (thus the pCRE in question are likely bound by TF(s) from a family that pCREs are similar to), the pairwise distances of known TF binding motifs (TFBM) across 30 TF families [50] were calculated and the 5^th^ percentile of distance, 0.39, (with a *p-value*=*0.05*) was set as a threshold [79].

To determine what pCREs identified in *S. lycopersicum* and *S. pennellii* for a given cluster are conserved across species (i.e. pCREs in question are likely bound by TF(s) of the same family), the pair-wise PCC (Pearson’s correlation coefficient) distance of the pCREs was generated with TAMO package [80] and used to construct the average linkage tree using UPGMA method in “cluster” package in R [81]. The threshold of 0.39 value that corresponds to the distance of the motifs among TF families were applied such that any pCREs within a branch length <0.39 are considered to be in the same sub-group and bound by TFs from the same family. Thus, the pCREs in a sub-group which contains pCREs from different spices are considered to be conserved between species (**Fig. 5A**, **Supplemental Fig. S4-S5, Supplemental Table S2**). The pCREs located in a given sub-group were merged through STAMP with default setting [82] to summary the sequence information of these pCREs since the pCREs are conserved but may be with subtle nucleotide difference and with various lengths (**Supplemental Fig. S6A-C**).

The similarity between the merged pCREs and known TFBMs (279 TFBMs of 256 plant TFs [50]) were determined with the threshold of PCC distance<0.39 (p<0.05) as described previously [79] (**Supplemental Fig. S6A-C**).

### Identification of orthologous genes

Using the longest protein sequences for genes, an all vs. all comparison of protein sequences was run on a combined set of genes in *S. lycopersicum* and *S. pennellii* using BLAST. Custom python scripts were then used to extract reciprocal best matches between species. The set of the reciprocal best matches was divided into those which were the best overall match (the “Overall” set) and those where one of the two proteins had a better match within species (the “Reciprocal-Only” set). Initially, there were 19,657 Overall and 1,198 Reciprocal-Only best matches. For Reciprocal-Only best matches, the sequence of the better within species matches were obtained, creating a group of 3 or more protein sequences (i.e. the best match between species gene pairs and any genes that are better matches within species) for each Reciprocal-Only best match.

For each pair of Overall best matches and group of Reciprocal-Only best matches, protein sequences were aligned using MAFFT. Protein alignments were then back-aligned to the longest coding sequences for genes in each species using custom python scripts. The resulting aligned nucleotide sequences were used to determine the Ks of best-matches using PAML. The “yn00” algorithm was used on sequence pairs from Overall best matches and the “codeml” algorithm was used for sequence groups from Reciprocal-Only best matches. Next, we visualized the distribution of Ks values for the “Overall” set because they have a clear 1:1 relationship between *S. lycopersicum* and *S. pennellii*. Given the recent speciation event, we expected the Ks distribution to follow a normal distribution. We observed a roughly normal distribution with a long right tail. We theorize that the extremely large Ks value in the tail can be attributed to ancient duplication events which experienced reciprocal loss in both species. Therefore, to enrich the set of reciprocal-best matches for orthologs of the recent speciation event, a normal distribution was fit to set of Ks values for the “Overall” set in R using non-linear minimization. The 99th percentile of the fit distribution was determined and applied as a cutoff to both the “Overall” and “Reciprocal-Only” best matches. This resulted in a final set of 16,222 orthologous genes between *S. lycopersicum* and *S. pennellii*.

### Gene ontology (GO) and metabolic pathway analyses

The datasets of GO annotation and metabolic pathways of genes in *S. lycopersicum* were retrieved from the Sol Genomics Network (https://solgenomics.net) and Plant Metabolic Network (http://www.plantcyc.org). To have comparable annotation set of GO and metabolic pathways of genes across species, the annotations of genes from *S. lycopersicum* were inferred to the orthologous ones in *S. pennellii*. In the end, 10,091 and 2,006 orthologous genes with biological process and metabolic pathway were retrieved for the downstream analyses. The list of orthologous gene pairs between *S. lycopersicum* and *S. pennellii* were generated as mentioned above.

The enrichment of GO terms and metabolic pathways in the clusters and differentially regulated gene sets, compared to the total orthologous genes, were determined though Fisher’s Exact test. A p-value obtained for each GO term and pathway comparison and was multiple-testing corrected [83].

### Conservation and divergence of pCRE sites in orthologous gene pairs

The region of the 1 kb upstream and 500 bp downstream of transcript start sites in the orthologous gene pairs were defined to be regulatory regions and aligned with MUSCLE package [84] (**Fig. 6A**). Based on the positions of the pCRE sites on the aligned sequences, these sites for each pCRE were assigned into 4 types: (1) “shared” (i.e. the site from each species was located on the same positions), (2) “specific” (i.e. the site was present only in one species), (3) “compensatory” (i.e. the site was present in both species but located in different location) and (4) “others” (i.e. any cases of pCRE sites were not assigned to the 3 types mentioned above). A likelihood score represent the conservation degree of pCRE sites for each pCRE was determined by taking the ratio of the pCRE site types (%) between the “shared” and “specific” ones. The orthologous gene pairs with consistent patter means the pairs are assigned to the same regulatory cluster as defined in **Fig. 2A;** otherwise, the OG pairs are considered to be with inconsistent patterns. Non-responsive orthologous genes are the orthologous genes if their fold-change values in any wound-treatment samples, compared to the control one, are between 1.2 and 0.8 in both *S. lycopersicum* and *S. pennellii* (n=1,371).

## DECLARATIONS

### Availability of data and material

The RNA-seq data from this study have been submitted to the Gene Expression Omnibus (GEO; http://www.ncbi.nlm.nih.gov/geo/) under accession numbers GSE93556.

### Authors' contributions

M.-J.L, G.A.H. and S.-H.S. designed the research. K.S. and G.A.H. performed the experiments. M.-J.L., S.U., N.P. and M.S.C. analyzed the data. M.-J.L., K.S., S.U., N.P., M.S.C., Y.M., G.A.H. and S.-H.S. wrote the paper.

## Acknowledgements

We thank for the members of the Shiu lab for discussion and Melissa Lehti-Shiu for helpful comments on the manuscript.

## Ethics approval and consent to participate

Not applicable.

## Consent for publication

Not applicable.

## SUPPLEMENTAL MATERIAL

**Supplemental Figure S1. Gene ontology (GO) category enrichment in wound-responsive genes**

(A) GO biological process categories enriched in wound up-regulated genes in different time point/location. Only the GO terms with false discovery rate adjusted p-value <1e-06 were shown. (B) As indicated in (A), but for wound down-regulated genes. Only the GO terms with adjusted *p*-value <1e-02 were shown.

**Supplemental Figure S2. Metabolic pathway enrichment in wound-responsive genes**

(A,B) Metabolic pathways significantly enriched in *S. lycopersicum* (*Sl*) or *S. pennellii* (*Sp*) genes from wound up-regulated clusters (A) and wound down-regulated clusters (B). (C,D) Pathways enriched in wound up-regulated genes (C) and wound down-regulated genes (D) in different time points/locations. Only the pathways with adjusted *p*-values <5e-02 were shown.

**Supplemental Figure S3. Genes differentially expressed between species prior to wounding**

(A) Heatmap showing differential expression where fold change (FC) values of all samples were calculated using the *S. lycopersicum* unwounded control expression values as the denominator. Only genes with significant FC values when comparing the unwounded controls between species are shown (n=826, |log_2_(FC)|>2 and adjusted *p*-values <0.05). (B) Differential expression values and test statistics by contrasting *S. pennellii* and *S. lycopersicum* unwounded controls of genes involved in biosynthesis of cuticular wax and cutin detected in this study and in the previous one [24].

**Supplemental Figure S4. Differential enrichment of pCREs in UUNN cluster genes between tomato species**

(A) Dendrogram showing the distances between the pCREs identified from UUNN cluster genes from in *S. lycopersicum* only (blue), *S. pennellii* only (orange), or both species (black). The dotted line indicates the threshold distance defined based on the 95^th^ percentile distances between binding motifs of TFs from distinct families and defines multiple pCRE sub-groups (numbered) where each sub-group contains pCREs likely bound by TFs of the same family (distance threshold=0.39). Subgroups with pCRE from only one species are labeled with asterisks. (B) Degrees of pCRE site enrichment in *S. lycopersicum* (blue) and *S. pennellii* (orange) UUNN genes. Adjusted *p*-value: multiple-testing corrected *p*-value. Dashed line, adjusted *p* <0.05. Yellow box: pCREs similar to the W-box (TTGAC[C/T]) [54]. The TGACG was more significantly enriched in *S. lycopersicum* whereas the ATGTCA (reverse complement, TGACAT) was more significantly enriched in *S. pennellii*.

**Supplemental Figure S5. Differential enrichment of pCREs in NDNN cluster genes between tomato species**

(A) Dendrogram showing the distances between the pCREs identified from NDNN cluster genes from in *S. lycopersicum* only (blue), *S. pennellii* only (orange), or both species (black). The dotted line indicates the threshold distance defined based on the 95^th^ percentile distances between binding motifs of TFs from distinct families and defines multiple pCRE sub-groups (numbered) where each sub-group contains pCREs likely bound by TFs of the same family (distance threshold=0.39). Subgroups with pCRE from only one species are labeled with asterisks. (B) Degrees of pCRE site enrichment in *S. lycopersicum* (blue) and *S. pennellii* (orange) NDNN genes. Adjusted *p*-value: multiple-testing corrected *p*-value. Dashed line, adjusted *p* <0.05.

**Supplemental Figure S6. Properties of the putative *cis*-elements regulating wound response**

(A,B,C) Correspondence (blue box) between the degenerate consensus sequences compiled from pCREs in a sub-group (rows) regulating genes in the UNNN (A), UUNN (B) and NDNN (C) cluster and TFs (columns, [50]) that likely bound the pCRE in question (see **Materials & Methods**). The matrices are consisted of pCREs that are conserved between both species (Conserved), specific to *S. lycopersicum* (Sl), or *S. pennellii* (Sp). Sequence logo: examples of compiled pCREs similar to the known defense responsive ABRE, G-box and W-box and the AT hook factors-bound elements discussed in the main text [34, 52-55, 66]. (D) Number of pCREs identified from one species that were (1) conserved and enriched in both species (gray, Type I), (2) conserved across species but specifically enriched in *S. lycopersicum* (cyan, Type II from *Sl*) or in *S. pennellii* (pink, Type II from *Sp*) and (3) nonconserved and specifically enriched in *S. lycopersicum* (light blue, Type III from *Sl*) or in *S. pennellii* (light pink, Type III from *Sp*) in three example wound response clusters. (E) Boxplot showing the wound response prediction performance (F-measure) based on a model using the pCRE sets in (D). For each wound response cluster, 10 F-measures were calculated from 10-fold cross validation and shown as a boxplot. Gray dot: the average F-measure of 10,000 random predictions indicating the performance of a meaningless model.

**Supplemental Figure S7. The types of pCRE sites on the aligned sequences of the orthologous gene pairs**

Box plots showing the proportion (%) of four types of pCRE sites found in orthologous genes with consistent regulatory patterns (Consistent, dark gray), inconsistent regulatory patterns (Inconsistent, unfilled), and the orthologous gene that are not responsive to wounding (Non-responsive OGs, light gray). The pCREs in UNNN (left), UUNN (middle) and NDNN (right) clusters were included.

**Supplemental Figure S8. MAKER-P-mediated improvement on S. pennellii gene annotation**

Cumulative Annotated Edit Distance (AED) plots showing the concordance of the available evidence with the gene annotation. An AED value of 0 to 1 indicates the complete absence to a perfect concordance of the evidences.

**Supplemental Figure S9. Similarity of gene expression profiles among replicates of samples**

Dendrograms showing the distance of the gene expression profile amongst samples and is determined by (1-spearman’s correlation). The *S. pennellii* W2hr_local_rep2 and W2hr_local_rep3 (red, bold) samples were not included for the downstream analyses since these 2 *S. pennellii* samples are more similar to *S. lycopersicum* samples. The *S. lycopersicum* W05hr_systemic_rep2 (black, bold) was also excluded for the downstream analyses since it showed poor correlation with other 2 replicates, *S. lycopersicum* W05hr_systemic_rep1 and W05hr_systemic_rep3. Red: *S. pennellii* samples; black: *S. lycopersicum* samples; W0.5hr and W2hr: 0.5- and 2-hr wound treatment; systemic_rep1-3: replicate1-3 from systemic leaves; local-rep1-3: replicate 1-3 from local leaves.

